# Breaking Medical Data Sharing Boundaries by Employing Artificial Radiographs

**DOI:** 10.1101/841619

**Authors:** Tianyu Han, Sven Nebelung, Christoph Haarburger, Nicolas Horst, Sebastian Reinartz, Dorit Merhof, Fabian Kiessling, Volkmar Schulz, Daniel Truhn

## Abstract

Artificial intelligence (AI) has the potential to change medicine fundamentally. Here, expert knowledge provided by AI can enhance diagnosis by comprehensive and user independent integration of multiple image features. Unfortunately, existing algorithms often stay behind expectations, as databases used for training are usually too small, incomplete, and heterogeneous in quality. Additionally, data protection constitutes a serious obstacle to data sharing. We propose to use generative models (GM) to produce high-resolution artificial radiographs, which are free of personal identifying information. Blinded analyses by computer vision and radiology experts proved the high similarity of artificial and real radiographs. The combination of multiple GM improves the performance of computer vision algorithms and the integration of artificial data into patient data repositories can compensate for underrepresented disease entities. Furthermore, the low computational effort of our method complies with existing IT infrastructure in hospitals and thus facilitates its dissemination. We envision that our approach could lead to scalable databases of anonymous medical images enabling standardized radiomic analyses at multiple sites.

## Introduction

The application of Artificial Intelligence (AI) in medicine promises to personalize diagnosis, decision management and therapy based on the combination of patient information with knowledge of thousands of experts and the outcome of billions of patient [1–4]. In recent years, a lot of scientific effort has focused on applications of AI in medicine with a particularly strong focus on radiology [5–10]. Whenever there has been progress towards this vision of an omniscient radiological AI, it has mostly been anticipated by corresponding technical advances in the field of Computer Vision on natural images. A very prominent example are convolutional neural networks (CNN), which had their breakthrough when more conventional computer vision algorithms were outperformed by a wide margin by AlexNet in 2012 [11]. Since then, CNNs have matched and even surpassed human performance on natural image recognition tasks [12]. Similar developments have taken place in medicine, where CNNs matched the performance of experts in CT-screening for lung cancer [13,14], retinal assessment [3,15] and intracranial hemorrhage detection [16]. However, human performance in computer vision on medical images was only achieved but not surpassed. Whenever human performance in computer vision on medical images has been reached, large datasets have been used - oftentimes pooled from many sites, containing thousands of images [5,17,18]. Going a step further and surpassing human performance in computer vision on natural images, however, required even larger databases containing up to billions of natural images [19].

Unfortunately, collecting and sharing such large amounts of medical images seems inconceivable at present, caused by their insufficient public availability and the lack of adequate data security. Even if the combined data worldwide would reach billions, like in the case of thoracic radiographs, patient privacy issues prohibit combining data from multiple sites. This is even more immediate given that the majority of patients are willing to share their data for research purposes, if adequate measures have been taken to protect their privacy [20]. Secure ways to share and merge medical image are very essential for the development of future computer vision algorithms [21] which underlines the urgent need for new concepts in this field. A very promising solution to overcome these limitations is the use of generative adversarial networks (GANs) which enable the generation of an anonymous and potentially infinite dataset of images based on a limited database of radiographs. GANs are a special class of neural networks that have first been introduced by Goodfellow [22] in 2014 and have since then been advanced to generate high resolution, photorealistic synthetic images [23]. By using training data, a GAN learns to generate new data that follows the same statistics leading to data (or images) that look as if they were real [23]. Very recently, GANs have been used successfully in the field of pathology to transform autofluorescence images into histologically stained versions of the same samples [24].

In this manuscript, we propose to use generative models based on GANs to break the boundary of sharing medical images and to enable merging of disparate databases without the limitations that are currently confining the collection of radiographs in a public database, see Figure 1. In order to demonstrate the performance of this new concept, we show that fully synthetic and thus anonymous images can be generated, which look deceivingly real - even to the expert’s eye - and that these images can be used in the medical data sharing process. Our concept is proposing how medical imaging or data can be shared in the near future and how a paradigm shift may pave the way to global databases of medical images akin to ImageNet, which revolutionized computer vision by providing “big data” to neural networks.

**Figure 1:**
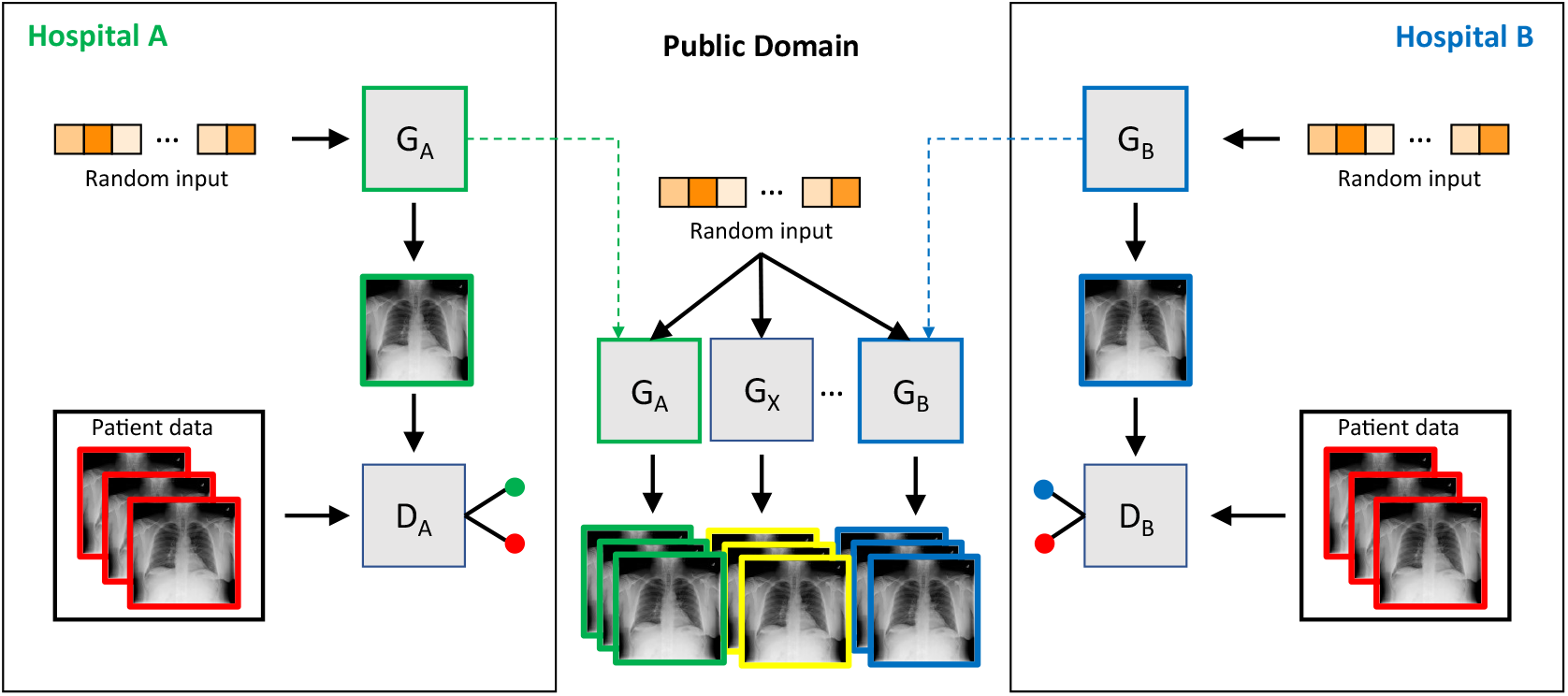
Concept of constructing a public database without disclosing patient sensitive data. The GAN in each hospital consists of a generator G and a discriminator D. During training, patient sensitive data (shown in red) is never exhibited to the generators directly. Only the discriminator gets to see patient sensitive data while trying to differentiate between real and artificial radiographs. After training is completed, only the generators - not the discriminators - are transferred to a public database and can be used to generate artificial radiographs. Thus, only the abstract information contained in the databases of contributing hospitals is transferred to the public domain.

## Results

### Generation of Artificial Radiographs

Generating artificial 2-dimensional images in high resolution is a non-trivial task and has just recently been made feasible by employing progressive growing during training [23] or by using large scale networks that require massive amounts of computing power [25]. As the computing power required for the latter approach is in general not accessible to most hospitals, we employed the first strategy in training our networks. Thus, the GAN was trained using progressively higher resolution stages starting with a resolution of 4×4 and stepping up in powers of 2 (8×8, 16× 16,…) to a resolution of 1024× 1024.

To demonstrate the feasibility of training a GAN at sites with limited access to computing power, we measured the time required for training with a publicly available dataset of 112,120 frontal radiographs [26] on a single Nvidia Tesla P100 graphics card: it took 60, 114 and 272 hours to train the GAN to generate radiographs with resolutions of 256×256, 512×512 and 1024× 1024, respectively. Once training had finished, inference, i.e. generation of artificial radiographs was much faster with a rate of 67,925, 41,379 and 4,511 generated radiographs per hour at the three resolution stages. Sample images are shown in Figure 2 (A)-(H) for a resolution of 256× 256. Further images for resolutions of 512× 512 and 1024 × 1024 are given in the supplemental Figure S2 and the supplemental Figure S3.

**Figure 2:**
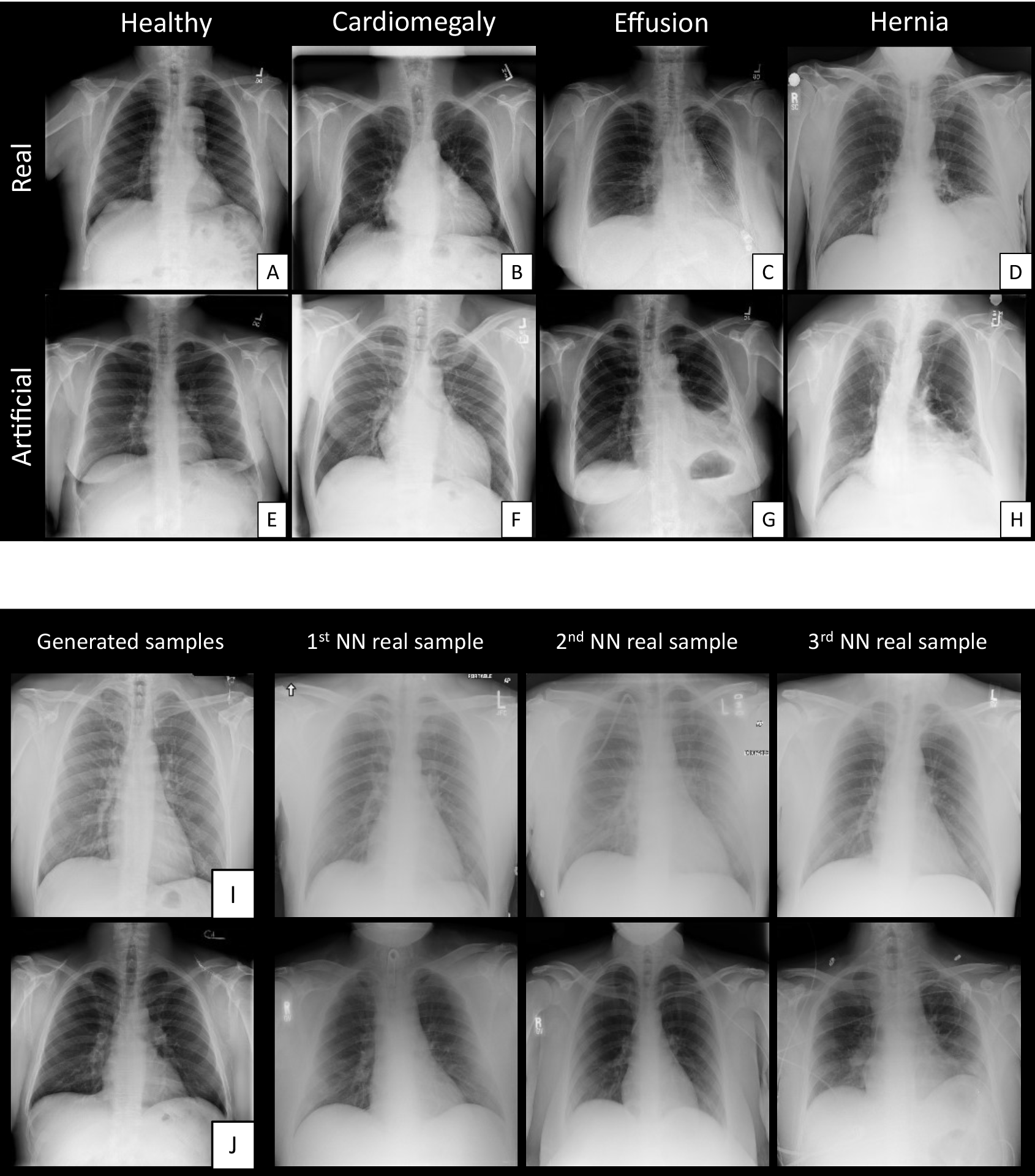
Comparison of real and GAN generated radiographs with a resolution of 256×256. Real thoracic radiographs from the database with labels healthy (A), cardiomegaly (B), effusion (C), and hernia (D) are displayed in the upper row. Artificial radiographs as generated by the GAN with the same labels are shown in the bottom row: healthy (E), cardiomegaly (F), effusion (G), and hernia (H). Note that consistent changes are seen in both the real and artificial radiographs: enlargement of the heart (A and F), opacification of the left lung (C and G) and mediastinal enlargement/shift (D and H). (I) and (J): randomly generated radiographs at a resolution of 256× 256 (left column) and their three nearest neighbors (NN) counterparts (i.e. most similar radiographs) in the real dataset in columns two to four. As implied by our network architecture, no duplication of an existing radiograph was found neither by visual inspection nor numerically by a high similarity measure.

### Ability of Human Readers to Distinguish Artificial Radiographs from real X-Ray Images

In order to test the quality of the artificial radiographs (i.e. radiographs generated by the GAN), six readers were each presented 50 artificial radiographs and 50 radiographs of real patients in randomized order, and the readers were separately tasked with deciding whether the presented radiograph was real or artificial. The tests were repeated with resolutions of 256 × 256, 512×512 and 1024 × 1024 amounting to a total of 18 tests with 100 radiographs each.

To assess whether experience with machine learning or radiological expertise is necessary to identify artificial radiographs, the readers were grouped and chosen as follows: group one consisted of three readers that had a background in computer vision (reader 1/2/3: 4/2/5 years of experience in computer vision respectively), while group two consisted of experienced radiologists (reader 4/5/6: 4/19/6 years of experience in radiology).

Accuracy in differentiating the GAN generated artificial images from the real images at resolutions of 256×256 was 60±5% for group 1 and 51±5% for group 2. Generating convincing radiographs at higher resolutions proved more difficult and experts were able to tell real from artificial radiographs more easily at resolutions of (512 × 512) / (1024×1024) with accuracies of 67±17% / 82±4% for group 1 and 65±5% / 77±13% for group 2. Thus, radiologists and computer vision experts performed similar when identifying artificial radiographs at high resolutions. While radiologists found errors more in anatomical details such as bone shape or rib cage morphology, computer vision experts tended to focus more on tiny details such as wave-like patterns, see the online supplemental figure S4. There was no inter-reader agreement between the readers for resolutions of 256, underlining the fact that identification of artificial radiographs at this resolution stage is hardly possible, see table 1. At higher resolutions, inter-reader agreement was consistently higher in accordance with the found higher accuracies in identifying artificial radiographs. It is important to note that these results were observed under restrictions: the radiographs were assessed on conventional 17 inch computer monitors without zooming into the images. The radiographs were presented in a given order: first the low-resolution radiographs, followed by the mid- and then the high-resolution radiographs. The readers were not allowed to go back and change previous decisions. When these restrictions were lifted, accuracy in determining whether a radiograph is real or fake was significantly increased: A radiologist with 9 years of experience who was given unlimited amounts of time and who first had a look at the high resolution radiographs on specialized radiological monitors to identify typical GAN related artefacts before going back to the 256× 256 radiographs was able to identify artificial radiographs in 86% of cases. Detailed results for each reader are given in the online supplemental table S3.

**Table 1:**
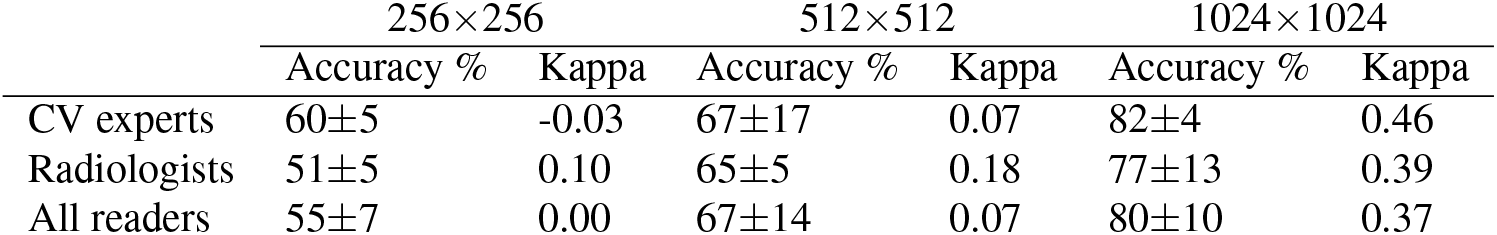
Real/artificial radiographs test. Accuracy and inter-reader agreement for the group of 3 computer vision (CV) experts, 3 radiologists and for all readers when deciding whether the presented radiograph is real or artificial, i.e. generated by a GAN.

|  | 256×256 |  | 512×512 |  | 1024×1024 |  |
| --- | --- | --- | --- | --- | --- | --- |
|  | Accuracy % | Kappa | Accuracy % | Kappa | Accuracy % | Kappa |
| CV experts | 60±5 | -0.03 | 67±17 | 0.07 | 82±4 | 0.46 |
| Radiologists | 51±5 | 0.10 | 65±5 | 0.18 | 77±13 | 0.39 |
| All readers | 55±7 | 0.00 | 67±14 | 0.07 | 80±10 | 0.37 |

The difficulties of the GAN to generate convincing radiographs at high resolutions are understandable, as the task becomes more difficult with the growing number of pixels: even for low resolution grayscale images of 100 × 100 pixels and 8 bit grey scale depth, the number of possible different images amounts to (10^4^)^256^, which is far greater than the estimated number of atoms in the universe. The GAN is tasked with identifying the subset of real-looking images out of this set that grows exponentially in size with increasing resolution. Not surprisingly, this process is not perfect and although the GAN manages to capture the general look of a real radiograph at high resolutions, small details reveal the artificial origin. After having performed the tests and with the knowledge of the ground truth, the raters conferred to identify these typical patterns that allowed for the differentiation of real from artificial images at high resolution. Among these were unphysiological configurations of the pulmonary vessels, aberrant bone structures and subtle periodic, wave-like patterns superimposed on the lung parenchyma, that reflect the network’s difficulty to generate fine details, see the supplemental Figure S4.

### Ensuring Retainment of Private Information

In order to exclude the possibility that the GAN memorizes and subsequently merely reproduces the given training examples, 300 randomly generated artificial radiographs were generated and their nearest neighbors in the database of real radiographs were sought according to the structured similarity index [27]. All 300 radiographs along with their respective 3 nearest neighbors were then plotted and a board-certified radiologist assessed whether an entity from the database of real radiographs had been duplicated.

In a set of 300 randomly drawn GAN images we did not find a single instance in which the artificial radiograph looked identical to its closest neighbor of the real dataset (Figure 2 (I) and (J)). In terms of the structured similarity index we did not find a single case in a set of 10^5^ randomly drawn artificial radiographs in which a digital twin was found in any of the real radiographs. More nearest neighbor comparisons can be found in the supplemental Figure S6.

It is plausible that the GAN does not duplicate images from the database of real radiographs as the generator will never have been in direct contact with a real patient radiograph. Only the discriminator of the GAN will have been presented real patient images. Transfer of the GAN discriminator network is, however, not needed in order to transfer the abstract information contained in the database as presented here – only the generator part of the GAN needs to be transferred. We envision, that this allows for an easy way of making the information contained within sensitive medical image databases publicly available: the sensitive and patient identifying information stays in-house, while the abstract information – more or less the radiological knowledge that a resident would have acquired if she or he had been trained on radiograph images – can be transferred as visualized in Figure 1.

### Performance of Classifiers Augmented with Anonymous Radiographs

We compared the performance of a state of the art classifier network that is trained on the full training set of original radiograph images versus the same classifier network when trained on only the GAN generated anonymized data. In the former case, the classifier reached an average AUC of 0.836±0.010% on the NIH test-set. Performing the same experiment with only the anonymous artificial data (n=78,468 radiographs) generated by the NIH-GAN, 90% of AUC, corresponding to a value of 0.750±0.010% was retained. We thus found that only employing the artificial radiographs from one institution allows for relevant information to be transferred to the classifier network without disclosing patient information, even though the achieved accuracies are lower in this single-institutional scenario. A detailed comparison of the AUC for each class and for both setups is shown in supplemental Figure 4A.

Nevertheless the GAN had difficulties in producing disease entities when one or more of the following conditions were met: 1) only few radiographs with that particular disease were present in the original dataset. 2) Disease location and appearance varied between different patients, e.g. the presence of nodules (can appear anywhere in the thorax) as opposed to cardiomegaly (always affecting the heart) and 3) accuracy of the labelling of the data of that particular disease was low (e.g. due to a somewhat fuzzy diagnostic boundary between infiltrations and pneumonia). Indeed, the quality of the ground truth labelling of the dataset has been criticized before [28, 29] and might be one of the main reasons for the reduced performance of training purely on the artificially generated radiographs. Another main reason is the limited size of the database of real radiographs that is available for training. Using more data - potentially even unlabelled data for pre-training of the GAN - will probably remedy these shortcomings. This is not out of reach considering that maximum care providers usually have databases of millions of radiographs, that they could use for the training of such GANs.

To simulate the data merging scenario as outlined in Figure 1, we compared the results of classification with a classifier solely trained on the NIH-GAN versus a classifier that was trained on both the NIH- and Stanford-GAN generated artificial images. Generated samples of our Stanford-GAN can be found in the supplemental Figure S5. The average values of the AUC, of accuracy, sensitivity, and specificity all increased significantly after integration of the artificial external dataset, see Figure 3. This demonstrates that the merging of multiple databases of artificially generated radiographs can further boost the performance of classifier networks and can indeed alleviate the performance bottleneck due to insufficient amounts of training data. Note that these performance improvements have been achieved without any techniques of domain adaption, i.e. without any efforts to homogenize the appearance of the radiographs of different databases. This not only offers opportunity for further performance improvements through domain adaption - currently an active area of research [30–32] - but also most likely makes the classifier network more robust to deployments in different environments, an important aspect in the translation of AI algorithms from workbench to clinics.

**Figure 3:**
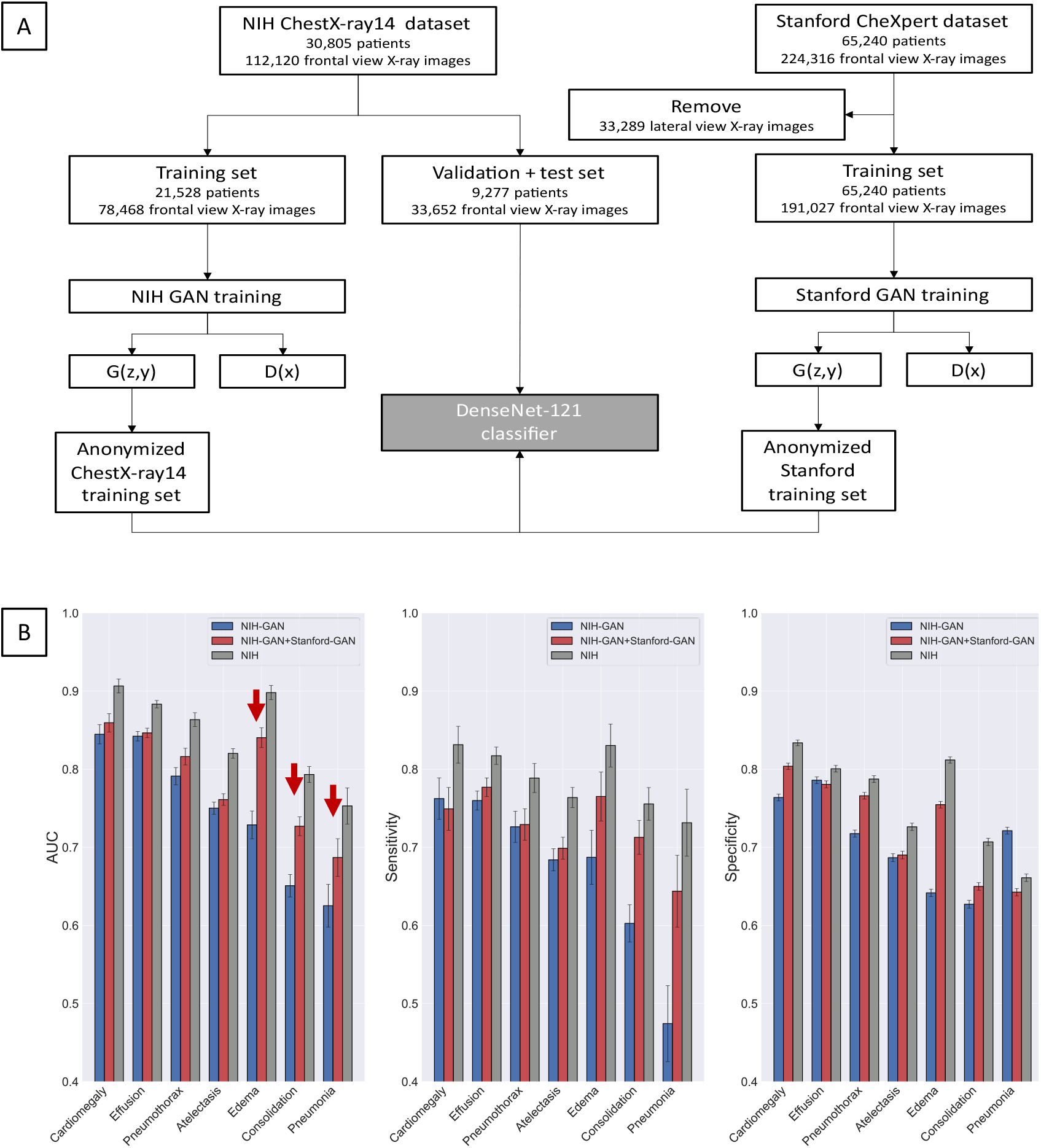
Using pooled artificial data from different sites, classification performance can be increased. In order to simulate the scenario in Figure 1, three classifiers were trained and compared: a classifier solely trained on anonymous radiographs generated with the NIH-GAN (blue), a classifier trained on the pooled anonymous dataset generated with the NIH-GAN and the Stanford-GAN (red) and a classifier trained on the non-anonymous NIH dataset with real radiographs (gray). The schematic of the data selection process is shown in A. AUC, Sensitivity and Specificity for the seven diseases are given in B. Note that especially the classification performances of formerly problematic cases such as edema, consolidation, and pneumonia were boosted by merging data from multiple sites (red arrows).

Finally, we investigated the effects of one class/pathology being heavily underrepresented in the training set: a specific hospital might have a large database of radiographs of one specific pathology, possibly due to its status as a specialized center, but might be lacking that of another pathology. To simulate this case we removed all but 5% of the cases with cardiomegaly from the training set and tested if a classifier can be trained with the conventional technique of oversampling [33], i.e. presenting the classifier with repetitive examples of the same radiograph multiple times during training. We found that this surprisingly led to a decrease in the AUC for that specific disease - possibly due to overfitting [34]. Even though overfitting might be avoided by careful tuning of the network and training process, a more generally applicable solution that led to better results without the need for manual intervention was the enrichment of that class with artificially created radiographs: while the AUC for cardiomegaly increased, we did not find any negative effects on the AUC of the remaining 13 pathologies, see Figure 4B.

**Figure 4:**
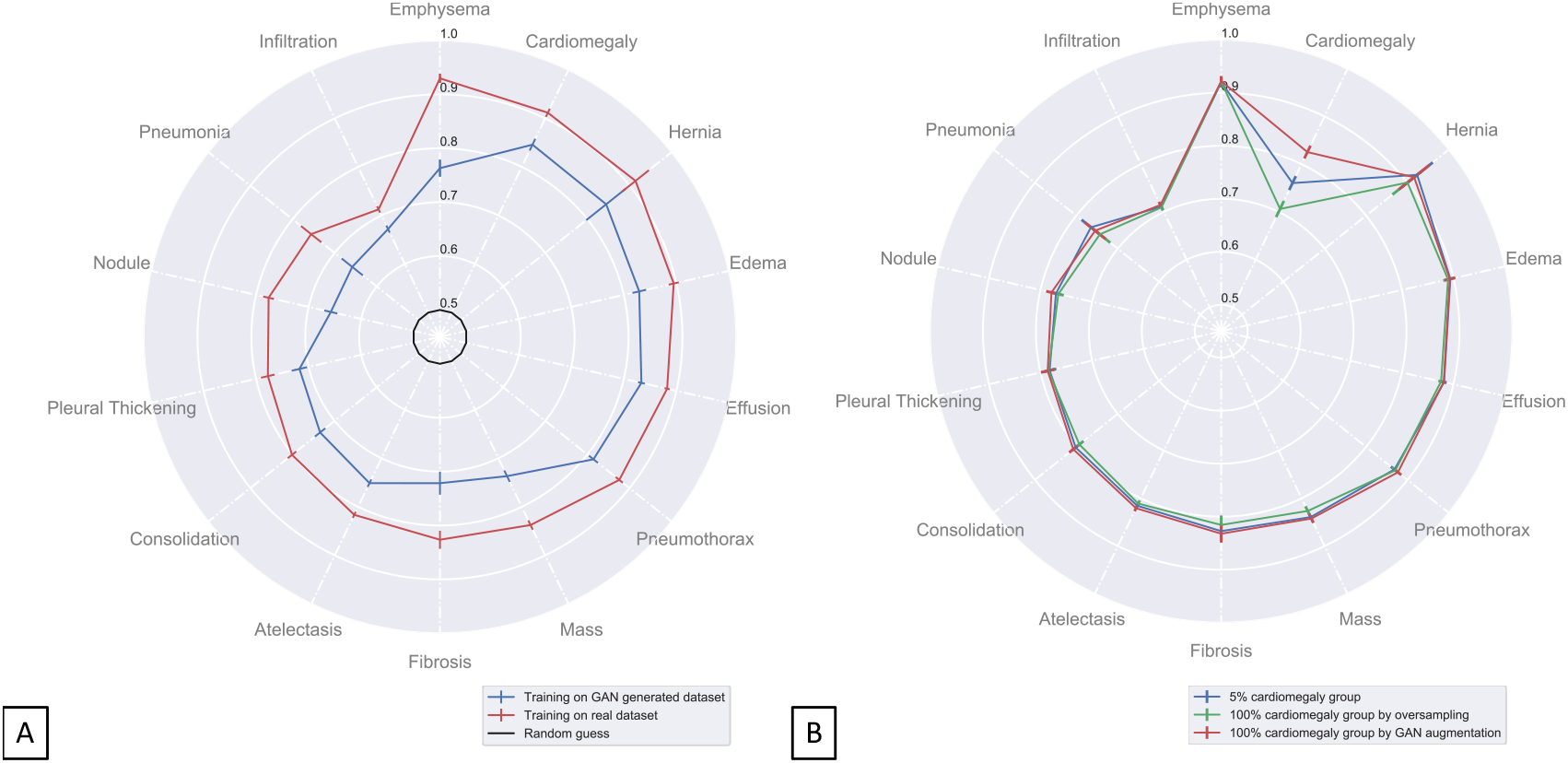
Classification performance of classifiers trained on anonymous datasets. (A): Class wise area under the ROC curve of classifier when exclusively trained on the GAN generated radiographs (blue) and when trained on the original dataset (red). A random guess amounts to an AUC of 0.5 which is shown in black as a reference. Standard deviations (shown as error bars) were obtained through bootstrapping. (B): Techniques for alleviating problems due to rare cases of one specific class (cardiomegaly). The training set was modified to only contain 5% of the original cardiomegaly cases. Blue curve: classifier trained with neither oversampling nor GAN augmentation resulting in a low AUC for cardiomegaly classification. Green curve: Oversampling of cardiomegaly decreased AUC, probably due to overfitting. Red curve: augmentation of the training set with GAN generated radiographs exhibiting cardiomegaly boosted the AUC of that specific class without affecting the other classes.

### Artificially Generated Images as a Visualization of what Neural Networks See

The images generated by the GAN can be specifically controlled: by changing the part of the input vector signifying the disease, radiographs with specific pathologies can be generated. We employ two techniques of visualizing the disease specific hallmark changes: first, the disease-specific entry in the input vector is gradually changed from zero to one while all other entries are kept at zero. The generated images then show the transition from healthy to diseased states and are stitched together to form an animation. Exemplary frames visualising the transition are given in Figure 5 for cardiomegaly and effusion. With cardiomegaly, we observe an enlargement of the projected heart shape reflecting the expected radiological change. Similarly, effusion shows the typical opacification of the lower lung field mirroring the collection of fluid there. Animations for all of the 14 disease states are given in the online supplement.

**Figure 5:**
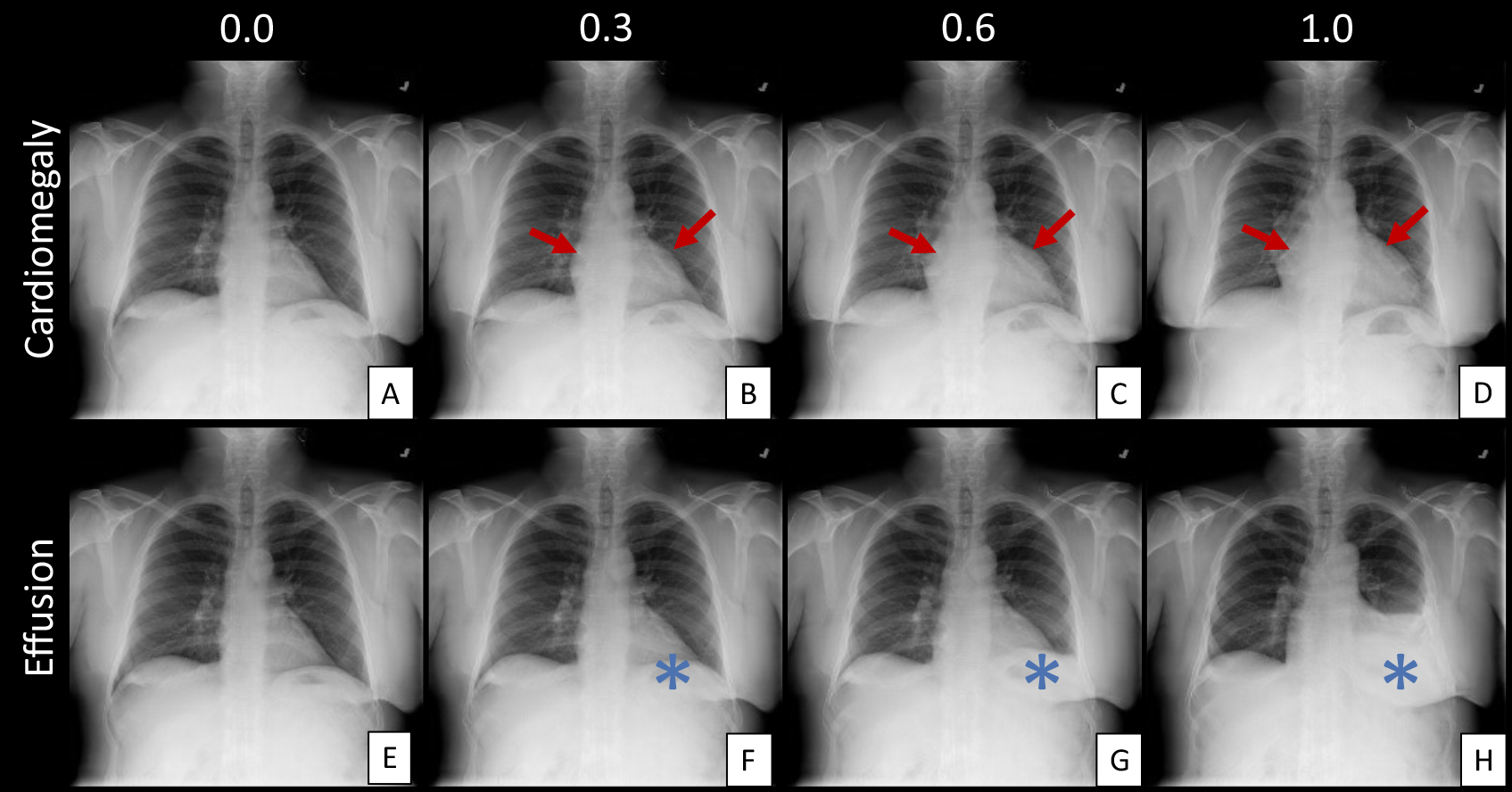
Quantitative pathology transitions. Transition from healthy to diseased GAN generated radiographs (top row: cardiomegaly, bottom row: effusion), when the latent vector that characterizes a specific pseudo-patient is held fixed and the entry in the one-hot encoding vector for that specific disease is increased from 0 (A and E), via 0.3 (B and F) and 0.6 (C and G) to 1.0 (D and H). Note that both cardiomegaly and effusion are realistically depicted: the heart shape grows from A-D, while the increasing opacification in E-F in the left lower lung reflects the accumulating amount of fluid which is less transparent for x-rays than lung tissue. Note that the left lung corresponds to the right image side as radiographs are depicted with the patient facing the reader. Cardiomagaly was marked by red arrows whereas effusion was marked by blue asterisks. Corresponding animations for both of these as well as the remaining 12 diseases are given as an online supplement.

Second, the pixelwise difference image between the fully diseased and the healthy radiograph is calculated and superimposed on the healthy radiograph as a colormap, see Figure 6 A for a schematic of the process. Examples of the such found visualizations are given in Figure 6 B for all of the 14 pathologies. One advantage of having full control over the disease state of the GAN radiographs is that any combination of diseases in a single radiograph can be generated by changing the corresponding entries in the input vector simultaneously. A possible clinical occurence is the combination of cardiomegaly with effusion. Artificial radiographs visualizing this transition in a two-dimensional array where the degree of cardiomegaly varies along one dimension and that of effusion along the other are given in the supplemental Figure S1. In accordance with the results found above, we found that the disease state as represented by the GAN transition is reflective of the underlying disease and agrees well with radiological expertise when many labelled examples of that disease were present in the original dataset and when the disease induced changes took place on a large scale of the radiograph (e.g. cardiomegaly or effusion) rather than on small patches at varying locations (nodules).

**Figure 6:**
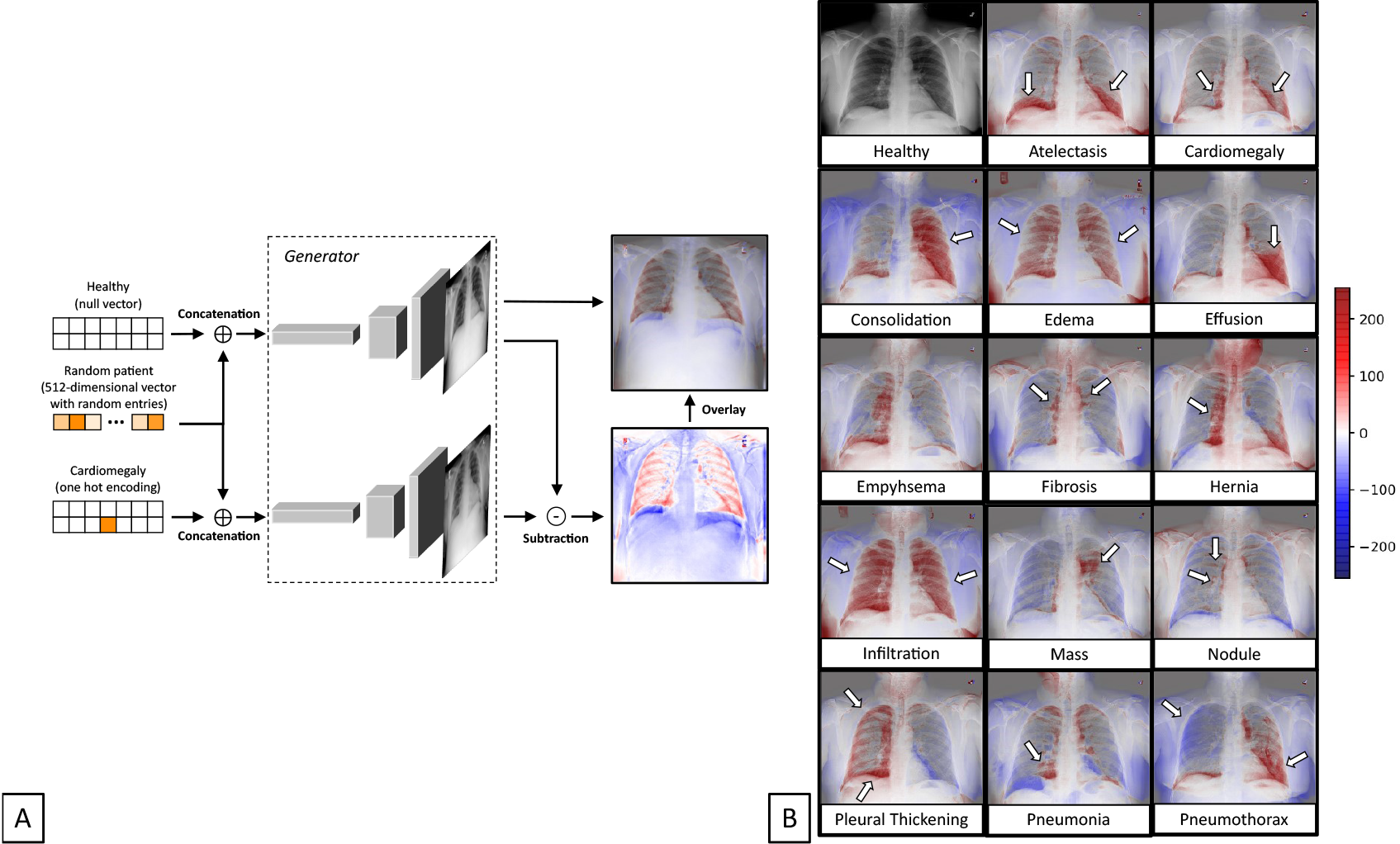
Visualizing learned pathological features. A: Generation of the disease specific pixel map: A randomly chosen vector with 512 Gaussian distributed entries characterizes one specific patient. The GAN is tasked with generating a healthy and a diseased radiograph of that patient (cardiomegaly in this example). A subtraction map is generated to denote the changes brought about by the disease and is superimposed as a color map over the generated healthy radiograph. B: Disease specific patterns generated by the GAN as shown in figure 6 for an exemplary randomly drawn pseudo-patient. Red denotes higher signal intensity in the pathological radiograph, while blue denotes lower signal intensity. Note that for some diseases such as cardiomegaly and edema, the pattern looks realistically, while the GAN struggles with diseases that have a variable appearance and where ground truth data is limited, e.g. pneumonia.

To uncover correlations between disparate pathologies, we let the classifier rate the score of a specific pathology when the GAN was tasked with generating an artificial radiograph of another disease and calculated Pearson’s correlation coefficient. We found that the clustering of related pathologies based on the correlation coefficient agreed well with clinical intuition: infiltration, pneumonia, consolidation, effusion and edema – all pathologies where at least one of the lungs opacifies - were related to each other, while being distant from e.g. emphysema or pneumothorax – diseases which are associated with increased radiolucency. The correlation coefficients for all 14 pathologies are given in Figure 7.

**Figure 7:**
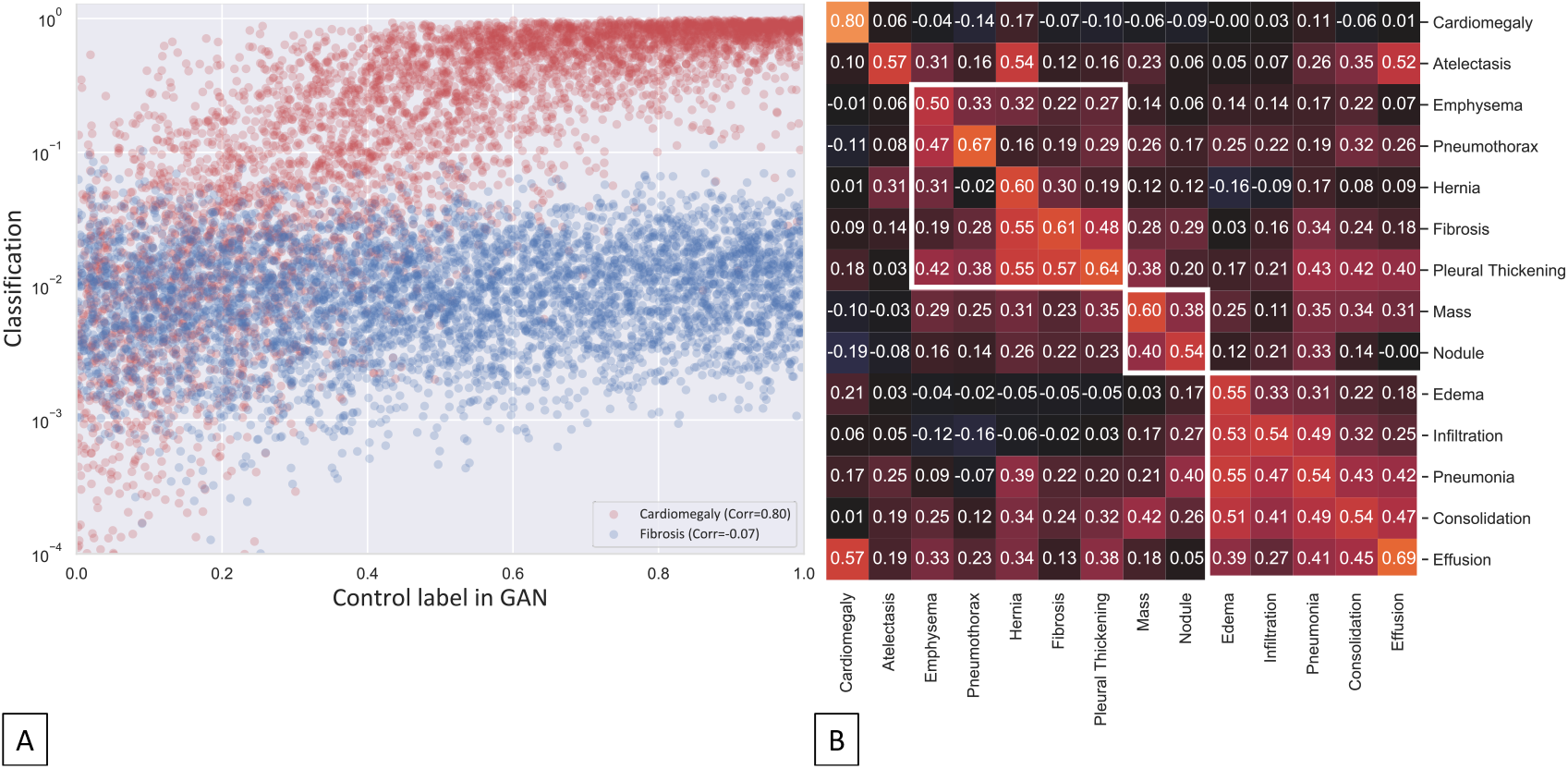
Revealing correlations within GAN generated pathological radiographs by the classifier trained on the real dataset. For each pathology, 5,000 random artificial radiographs with a pathology label drawn from a uniform distribution between 0.0 and 1.0 were generated. The images were then rated by the classifier network and Pearson’s correlation coefficient was calculated for each pairing of pathologies (shown in A with the GAN cardiomegaly label on the x-axis and the cardiomegaly and fibrosis classifier output on the y-axis in red and blue respectively). Resulting correlation coefficients for all 14× 14 pairings are displayed and color-coded in B. Clustering on the x-axis (i.e. the GAN label axis) was performed in order to group related diseases together. The obtained clustered blocks are marked with white bordered boxes.

## Discussion

In this study we demonstrated that GANs can be used to generate deceivingly real looking radiographs and to merge databases of radiological images without disclosing patient sensitive information. This contributes to solving the important problem of building large radiological image databases for the training of computer vision algorithms. While radiographs may in principle be abundantly available, universal access is in general severely restricted due to data protection laws: privacy concerns restrict the export of sensible patient information to extramural institutions and often only a small fraction of the available data can be used (e.g. patient consent form not universally available). In these cases, GANs that have been trained in-house may serve as a mean to distribute the information contained within the database without actually providing a real snapshot of patient sensitive data: only the weight distribution of the GAN needs to be transferred and a representative artificial dataset of millions of radiographs may be generated in reasonable computational time at a peripheral site. This is in contrast to previous works of Shin et al. in which lower resolution artificial images could be produced but always required a recourse to the original patient images as inputs to an image to image translational network [35, 36]. We demonstrate the feasibility of using GANs as a tool of effective oversampling when the pathology distribution within a medical dataset is highly imbalanced. Moreover, our developed GANs can be used to visualize what the generator neural network sees and to reveal correlations between diseases. This might be used in several ways: as a check that the GAN has been trained correctly, as a tool to uncover relationships between diseases or to visualize hallmark changes of pathologies [37] and potentially also as a decision support system for diagnosis.

Another advantage of the proposed concept of data sharing is the immense reduction in data storage requirements: storing of a single radiograph image with a resolution of 1024 × 1024 and a bit depth of 8 bits requires 1 megabyte of hard drive space. Considering that the number of radiographs acquired at a maximum care hospital easily reaches 100,000 per year, constructing a database of millions of radiographs (ImageNet comprises e.g. 14 million natural images) is not out of reach. While the actual images themselves were to take up terabytes of data, the storing of the weights of the GAN generators only amounts to a few hundred megabytes (in our case about 23 million weights with a precision of 32 bits: 80 MB of storage). The GANs thus also serve as a data content compression tool that might promote the development of computer vision algorithms at smaller institutions that do not have access to large data storage capabilities.

The concepts demonstrated in this work rely on two-dimensional images, but there is no principal restriction on the number of dimensions that the real and artificial images are allowed to have, or even that the data has to consist of images only [38]. Thus, the same concept could be translated to volumetric computed tomography or magnetic resonance images, to fluoroscopy, to time-series of volumetric data (e.g. contrast-enhanced computed tomography), or even to imaging data in conjunction with clinical data (e.g. an MRI with associated expression profiles of laboratory tumor markers). However, due to the exponentially increasing size of the data space, we expect that the problem of generating artificial data of very high-dimensionality is much more difficult and that a far greater number of real cases would be needed for the GAN to converge.

We hope that the work presented in this manuscript makes a leap towards the goal of a medically omniscient AI that provides expert knowledge to everyone at anytime by making “big data” available to the worldwide research community.

## Materials and Methods

### Dataset and Preprocessing

Two datasets were used in this study: first, the ChestX-ray dataset released by the NIH in 2017, containing 112,120 frontal radiographs of 30,805 unique patients [10]. At the time of its publication this dataset comprised 8 disease entities and was later updated to contain 14 pathologies [26]. We employ the full set of 14 pathologies in this study. To ensure that no information leaked into the test-set used for the evaluation of the algorithms, patient-wise stratification into training- (21.528 patients, 78.468 radiographs, 70%), validation- (3.090 patients, 11.219 radiographs, 10%) and test-set (6187 patients, 22.433 radiographs, 20%) was performed before any further steps and the test-set was kept separately until the final testing of the algorithms. A detailed label statistics for the ChestX-ray 14 dataset is given in the supplemental Section 1 and the supplemental Table S2.

The second dataset used in this study is the CheXpert dataset, which has been released by Irvin et al. in January 2019 and contains 224,316 chest radiographs of 65,240 patients [39]. This dataset was used to train a second GAN in order to demonstrate the feasibility of the proposed data sharing approach (see Figure 1). A detailed explanation of the label preparation and statistics for the CheXpert dataset is given in the supplemental Table S2. Testing of all classification algorithms was done on the NIH test-set. Therefore, no subdivision of the CheXpert dataset into test, training and validation sets was needed and all available frontal radiographs of the CheXpert dataset (n=191,027) were used for training of the GAN.

The radiographs’ intensity values were normalized to the range 0-255 and image resolution was resized such that each radiograph was available at resolutions of 256×256, 512×512 and 1024× 1024.

### Model Architecture and Implementation

Two neural network architectures were used in this paper: Firstly, generative adversarial networks (GAN) as introduced by Goodfellow [22] were adapted to incorporate an input condition [40] to selectively generate artificial radiographs with a certain pathology. To generate high resolution images we employed progressive growing, a technique in which the GAN is trained in progressively higher resolution stages [23]. The used GAN architecture and training details can be found in the supplemental Section 2. Secondly, a densely connected convolutional neural network - with 121 layers was used as a classifier. It was pretrained on 14 million natural images (ImageNet database [11]) and subsequently trained on the radiographs in this study. This architecture has been shown to achieve state of the art performance on the ChestX-ray dataset [41] [42] before. Implementations were done using TensorFlow 1.9.0 [43] and PyTorch 1.1.0 [44].

### Training of the GANs

We trained two GANs based on two separate datasets: on the NIH training subset and on the Stanford radiographs in a progressive growing strategy. Training proceeded in repetitive stages: once training of one resolution stage stabilized after being presented 600,000 real radiographs, the layers responsible for the next resolution stage were gradually faded in and training continued with another 600,000 radiographs during this fade-in stage. In total the GANs were each presented 12 million radiographs and thus learned to first explore the large scale pattern and overall contrast before focusing their attention towards finer details.

### Training of the Classifier with Real and Artificial Data

All classifier models utilized validation-based early stopping with sigmoid binary cross entropy loss as the criterion. No oversampling of underrepresented classes was employed except for the experiment in which we specifically tested for the effect of oversampling. Training of the classifier network was done for a variety of different settings.

The settings were as follows: Firstly, the classifier was trained on the full training and validation set of real images. Secondly, the NIH-GAN was used to generate the same number of radiographs as in the training and validation sets and the classifier was trained on these sets.

In another experiment, we evaluated how the inclusion of external datasets as laid out in Figure 1 can improve the classification performance. For this we trained the classifier on artificial radiographs produced by both the NIH-GAN and the Stanford-GAN. As not all of the 14 pathologies labelled in the NIH dataset had been labelled in the CheXpert dataset, we only classified those pathologies that were present in both datasets’ labels, namely cardiomegaly, effusion, pneumothorax, atelectasis, edema, consolidation and pneumonia.

Finally, we conducted an experiment in which GAN generated images were rated by the trained classifier network to discover correlations between disparate pathologies. For each pairing of pathology as generated by the GAN and pathology as classified by the classifier, we calculated Pearson’s correlation coefficient and performed clustering on the resulting correlation matrix.

### Reader Study

Six readers were tasked with identifying whether a radiograph was real or fake. The tests were performed as follows: Each reader was given a time span of 30 seconds within she/he had to decide whether the presented radiograph was real or fake. To prevent readers from identifying GAN related features on the high resolution radiographs first - which are harder to produce and thus presumable more prone to artefacts - and transferring that knowledge to the low resolution images, the radiographs were presented in the following order: 100 radiographs of 256 × 256, 100 radiographs of 512×512 and finally 100 radiographs of 1024×1024. All presented radiographs were different, i.e. the 256 × 256 were different from the 512×512 and from the 1024 × 1024 radiographs. Reading tests were done on a 17 inch computer monitor.

### Statistical Analysis

For each of the above described experiments we calculated the following parameters on the test set: area under the curve (AUC), accuracy, sensitivity and specificity. To assess the errors due to sampling of our specific test set we employed bootstrapping with 10,000 redraws. An exemplary bootstrapping analysis of Figure 3 can be found in the supplemental Figure S7. The standard error of the accuracy in the real vs. artificial tests for each human reader was calculated among the reader performances and Fleiss’ kappa was employed to assess inter-reader agreement between readers.

## Data availability

This study used two publicly available dataset: NIH ChestX-ray14 dataset: https://nihcc.app.box.com/v/ChestXray-NIHCC; Stanford CheXpert dataset: https://stanfordmlgroup.github.io/competitions/chexpert/. The full images used in our real/artificial radiographs test are available on https://drive.google.com/open?id=1_snb7hQ47WYxJEYK95G3cYlWSqckvRDW. Images and animations to demonstrate controlled pathology transitions can be found under https://drive.google.com/open?id=1rBni9KSn1iB0OMALz8XJHDQgZqiBfgA-.

## Code Availability

Details of the implementation as well as the weights of the neural networks after training and the full code producing the results of this paper are made publicly available under https://github.com/peterhan91/Throax_GAN.git.

## Acknowledgments

This research project was supported by the START program of the Faculty of Medicine, RWTH Aachen, Germany, through the START rotation program granted to D.T. and by the DFG, Germany through a grant given to S.N.

## Author contributions

T.H., D.T., V.S., and F.K. conceived the idea and approach. F.K., V.S., S.R., S.N., C.H., N.H., D.M., and D.T. contributed to the experiments. T.H., D.T., C.H., and N.H. developed the code infrastructure and GAN training setup. T.H., D.T., F.K. and V.S. wrote the manuscript.

## Online supplement

### 1 Preprocessing steps in CheXpert dataset

#### Frontal radiographs filtering

The Chexpert dataset consists of both frontal (191,027) radiographs and lateral radiographs. Only frontal thoracic radiographs were used for the training of the Stanford-GAN.

#### Generating labels

1. Pathology labels not mentioned (NaN) were assigend to 0.0.
2. According to the labeling performance comparison mentioned in [39], most of the uncertainty labels (U) were assigned to 1.0, except for the consolidation class.
3. Uncertainty labels for consolidation were assigned to 0.0.
4. A Python dictionary was used to record pathology labels of radiographs: the key was set to the radiograph filename and the corresponding value was set to the one-hot pathology vector.

#### Preprocessing of Radiographs

The aspect radio of radiographs was rescaled to 1:1 by zero-padding. The pixel intensity of all radiographs was normalized to the range of 0 to 255.

### 2 Network training details

We used mirrored structures for our generator and discriminator in the GAN and they always grow in synchrony. The training starts with a 4×4 resolution and continues until the discriminator has been shown 15 million real images in total. Both in the generator and the discriminator we used leaky ReLU with leakiness of 0.2 as the activation function. The loss function of our GAN was defined as the Wasserstein loss with gradient penalty [45]. In addition, the batch size of GAN training was tailored for a high-end Nvidia GPU with 16 GB of RAM:

1. For resolution stage 4×4 to 32×32, we employed a batch size of 128.
2. For resolution stage 64× 64 and higher, the batch size was gradually decayed by a factor of 2 to ensure that the batch fits into the memory: 64×64 → 64; 128×128 → 32; 256×256 → 16; 512×512 → 8; 1024×1024 → 4.

**Table S2:** Data characteristics of NIH ChestX-ray 14 and Stanford CheXpert dataset.

|  | NIH ChestX-ray 14 dataset | Stanford CheXpert dataset |
| --- | --- | --- |
| Number of patient radiographs | 112,120 | 224,316 |
| Number of patients | 30,805 | 65,240 |
| Age, mean (SD), years | 46.9 (16.6) | 60.7 (18.4)* |
| Number of females (%) | 48,780 (43.5%) | 90,778 (40.6%)* |
| Number of pathology labels | 14 | 14 |
| Cardiomegaly | 2,776 | 30,092 |
| Edema | 2,303 | 61,493 |
| Effusion | 13,317 | 86,477 |
| Pneumothorax | 5,302 | 20,401 |
| Atelectasis | 11,559 | 59,583 |
| Consolidation | 4,667 | 12,983 |
| Pneumonia | 1,431 | 20,656 |

Note. – \*Patient data for the 223,414 / 224,316 radiographs in Stanford CheXpert dataset.
The seven overlapping labels between the NIH and Stanford dataset are listed in the table above. Note that pathological radiographs seem to be more common in the Stanford CheXpert dataset due to the differences in the labelling process [39].
Abbreviations: NIH, National Institutes of Health Clinical Center; SD, standard deviation.

**Table S3:** Performance measures of participants in the GAN real/fake test. Note that the reading conditions for Radiologist (RAD) 4 were different from the remaining readers: Radiologist 4 performed the test at a dedicated radiological working station and was allowed to first analyse the high resolutions (1024× 1024) radiographs for telltale signs of the GAN before going back to the low resolution examples.

|  | 256 × 256 |  |  | 512×512 |  |  | 1024×1024 |  |  |
| --- | --- | --- | --- | --- | --- | --- | --- | --- | --- |
|  | Accu. % | Sens. % | Spec. % | Accu. % | Sens. % | Spec. % | Accu. % | Sens. % | Spec. % |
| CV 1 | 53 | 52 | 54 | 90 | 94 | 86 | 86 | 88 | 84 |
| CV 2 | 62 | 70 | 54 | 62 | 74 | 50 | 77 | 80 | 74 |
| CV 3 | 66 | 76 | 56 | 51 | 78 | 24 | 84 | 96 | 72 |
| Rad 1 | 45 | 50 | 40 | 66 | 66 | 66 | 69 | 74 | 64 |
| Rad 2 | 58 | 66 | 50 | 53 | 68 | 38 | 68 | 76 | 60 |
| Rad 3 | 50 | 48 | 52 | 77 | 90 | 64 | 96 | 98 | 94 |
| Rad 4 | 86 | 76 | 96 | 96 | 92 | 100 | 100 | 100 | 100 |

Abbreviations: CV, computer vision experts; Rad, radiologists with more than 5 years experiences; Aver., average; Accu., accuracy; Sens., sensitivity; Spec., specificity.

**Figure S1:**
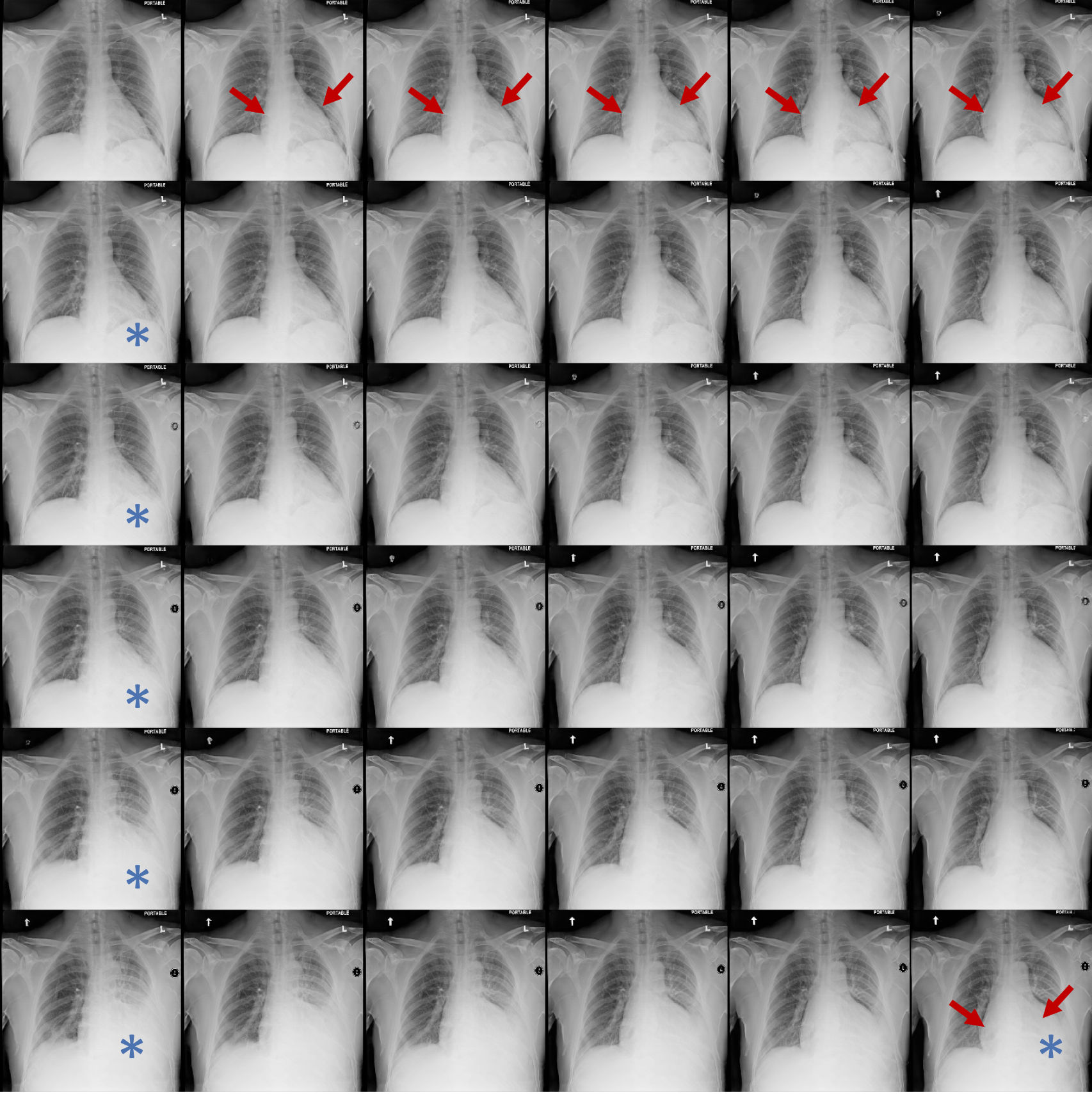
Generation of artificial radiographs with multiple pathologies. Snapshot of radiographs that have been conditioned to show both cardiomegaly and effusion. Along the horizontal direction, the cardiomegaly label was conditionally changed from 0.0 to 1.0 with a stepsize of 0.2. Similarly, the effusion label was conditionally changed from 0.0 to 1.0 with a stepsize of 0.2 along the vertical direction. Cardiomagaly was marked by red arrows whereas effusion was marked by blue asterisks.

**Figure S2:**
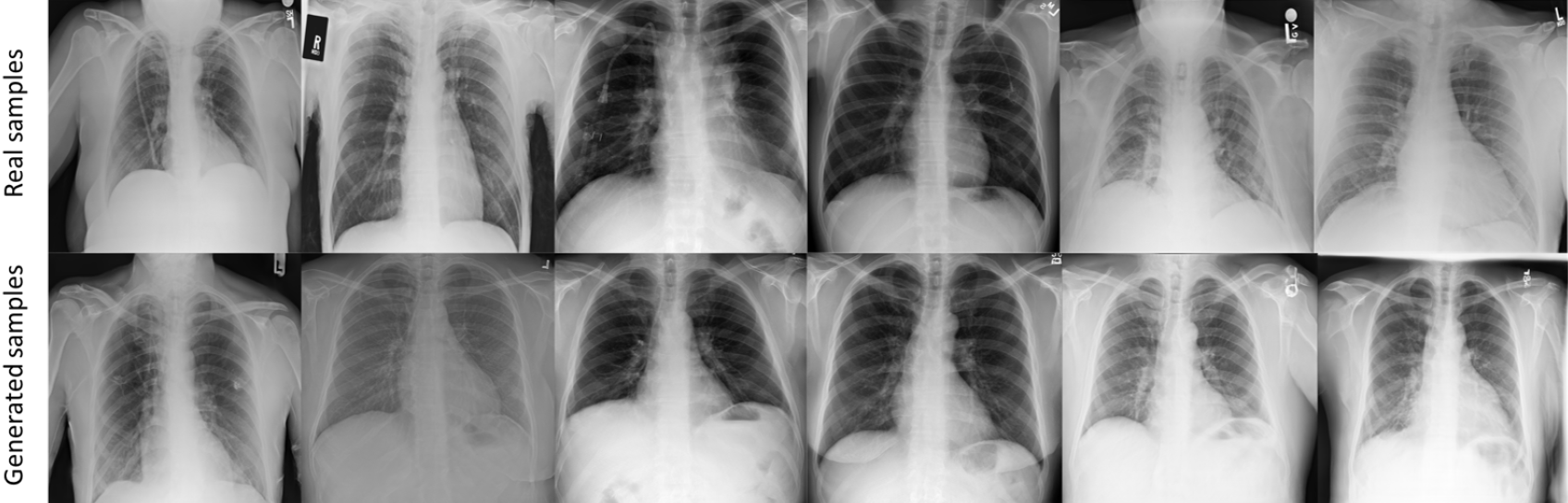
Sample images used for the test of distinguishing real and NIH-GAN generated radiographs with a resolution of 512 × 512.

**Figure S3:**
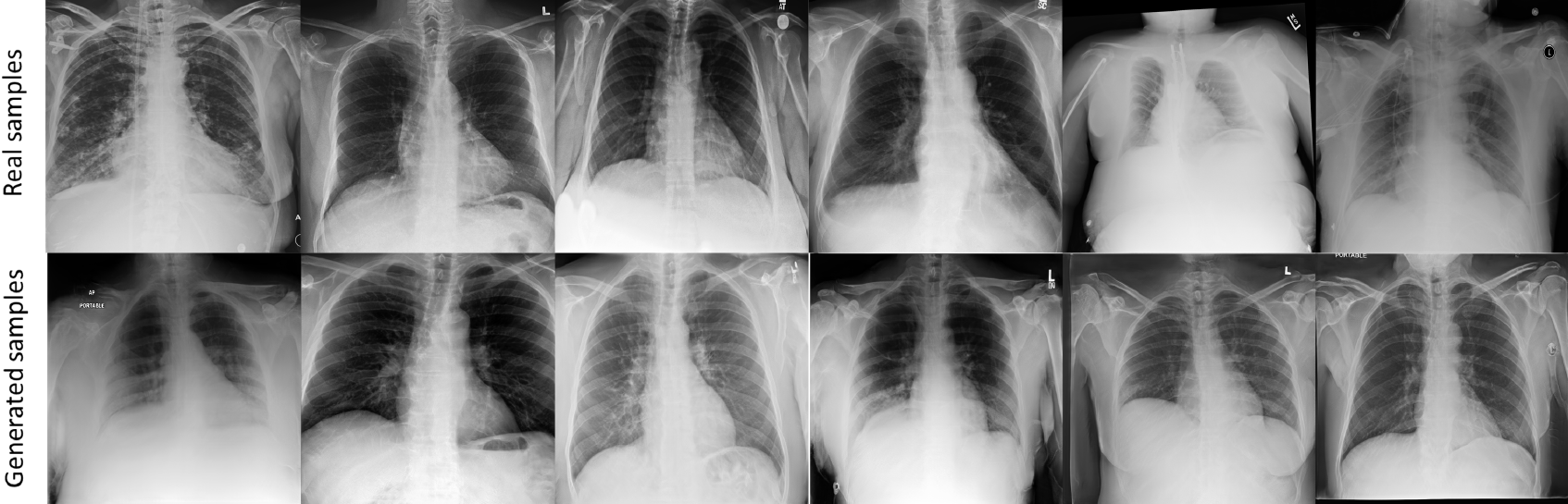
Sample images used for the test of distinguishing real and NIH-GAN generated radiographs with a resolution of 1024 × 1024.

**Figure S4:**
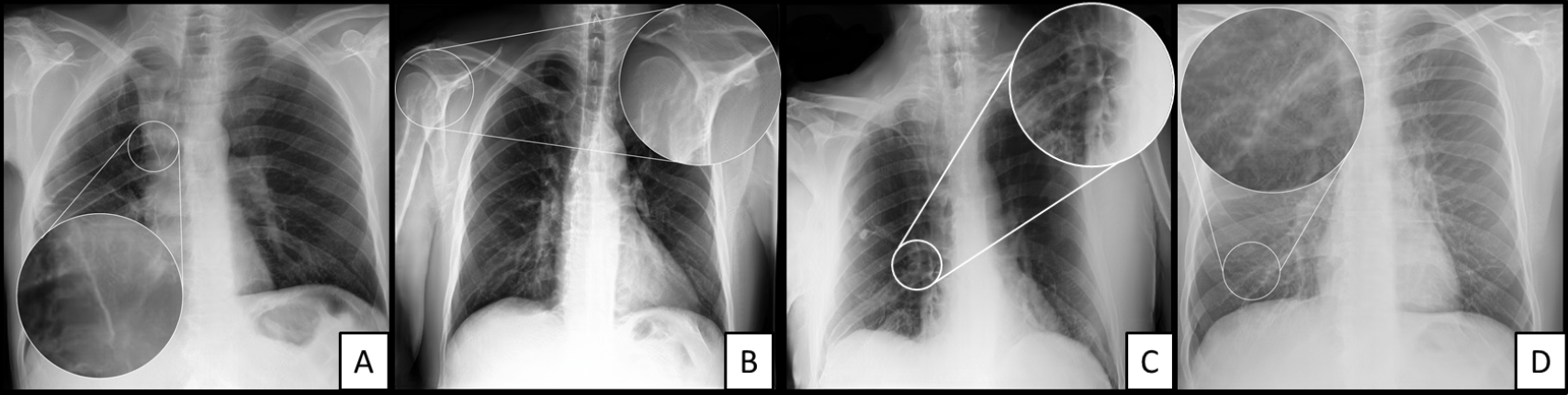
Artifacts in artificial radiographs with resolution 1024 × 1024. GAN generated high-resolution images with a resolution of 1024 × 024 look real from far away. However, details reveal the artificial origin of the data. From left to right: dense line in unphysiological orientation adjacent to mediastinum (A), bizarre configuration of the humeral head (B), unphysiological configuration of right pulmonary vessels (C) and periodic, wavelike pattern superimposed on the lung parenchyma(D). Note, that those artefacts do not appear consistently and even though the GAN has difficulties reproducing all details of an radiograph collectively, it often generates them in a realistic matter: the humeral head (artefact B) is correctly depicted in C, while the aberrant vessel structure of the right hilus (artefact C) is correctly depicted in B.

**Figure S5:**
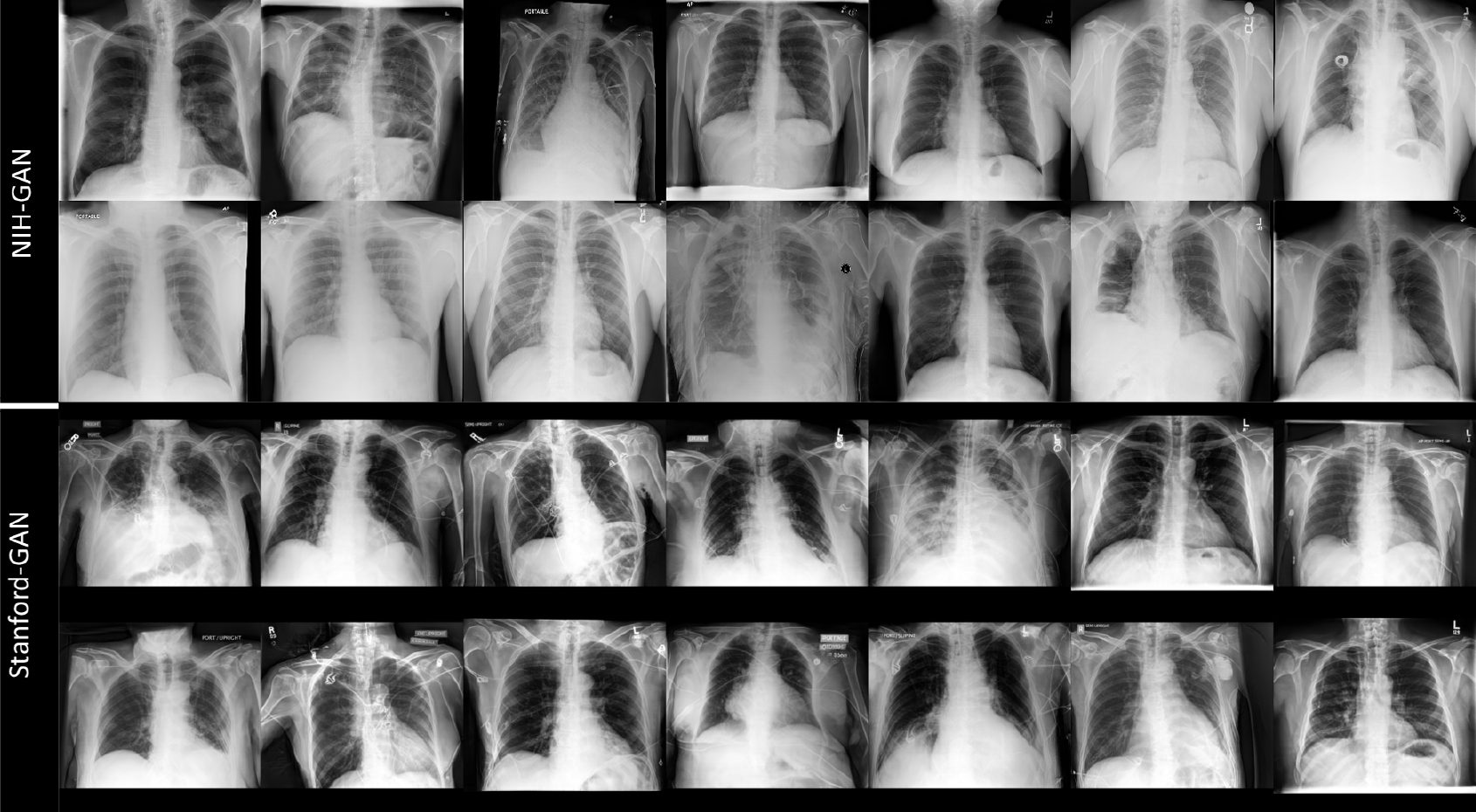
Comparison of NIH-GAN and Stanford-GAN generated radiographs with a resolution of 256 × 256. Both image contrast and aspect ratios vary between the NIH-GAN and the Stanford-GAN generated samples. These differences are due to existing differences in the database of real radiographs, probably because of different standards of how the radiographs were acquired. In our manuscript, we did not perform any domain adaption techniques, thus, these differences offer room for performance improvement in the future when applying such techniques in the proposed anonymous data merging experiment.

**Figure S6:**
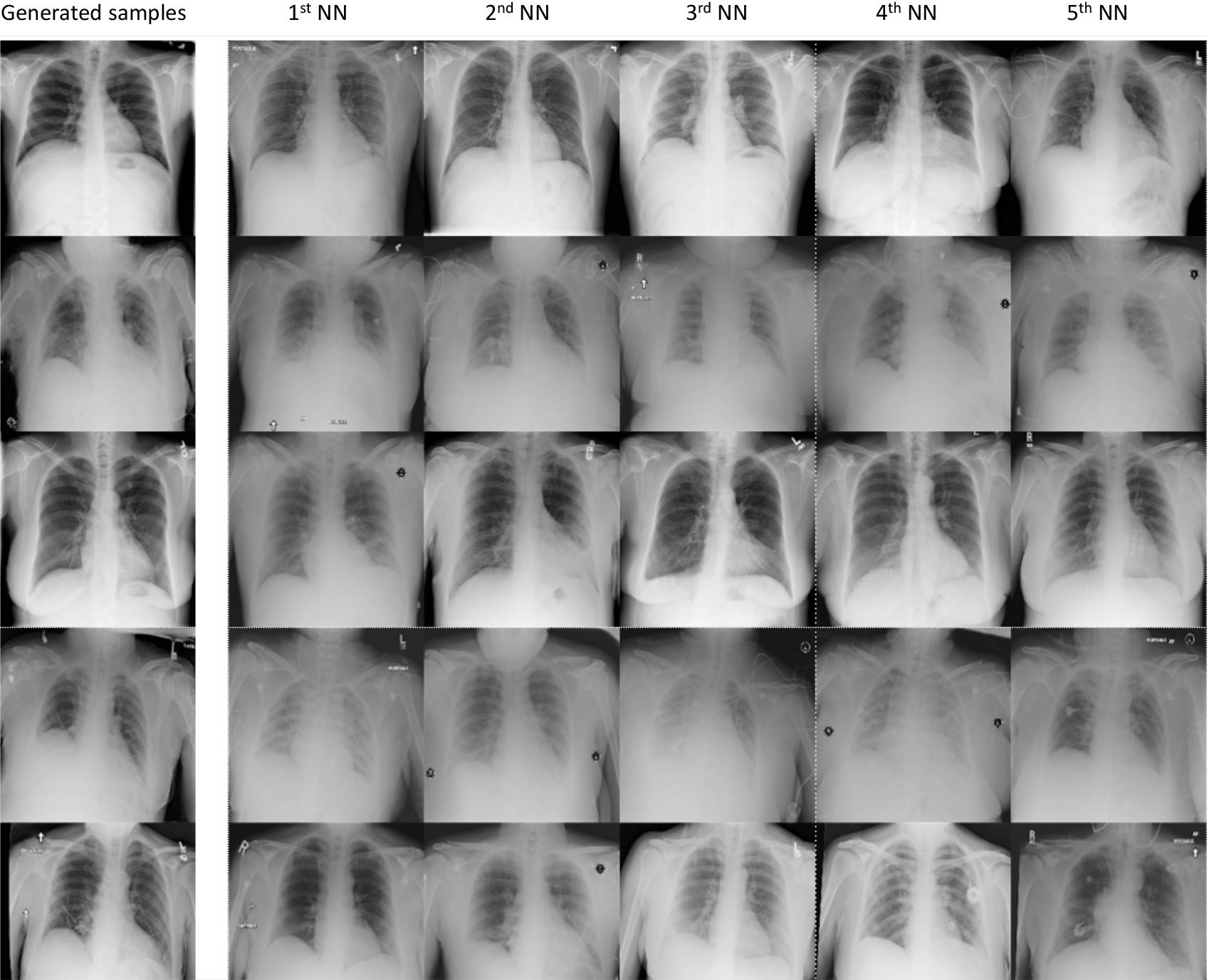
Randomly generated radiographs by our trained generator network at a resolution of 256 ×256 (leftmost column) and their five nearest neighbors (NN) counterparts in the real dataset (columns 2-6). No duplication of an existing radiograph was found neither by visual inspection nor numerically by a high similarity measure. Patient sensitive information can be protected during the data communcation process by transfering only the trained generator network.

**Figure S7:**
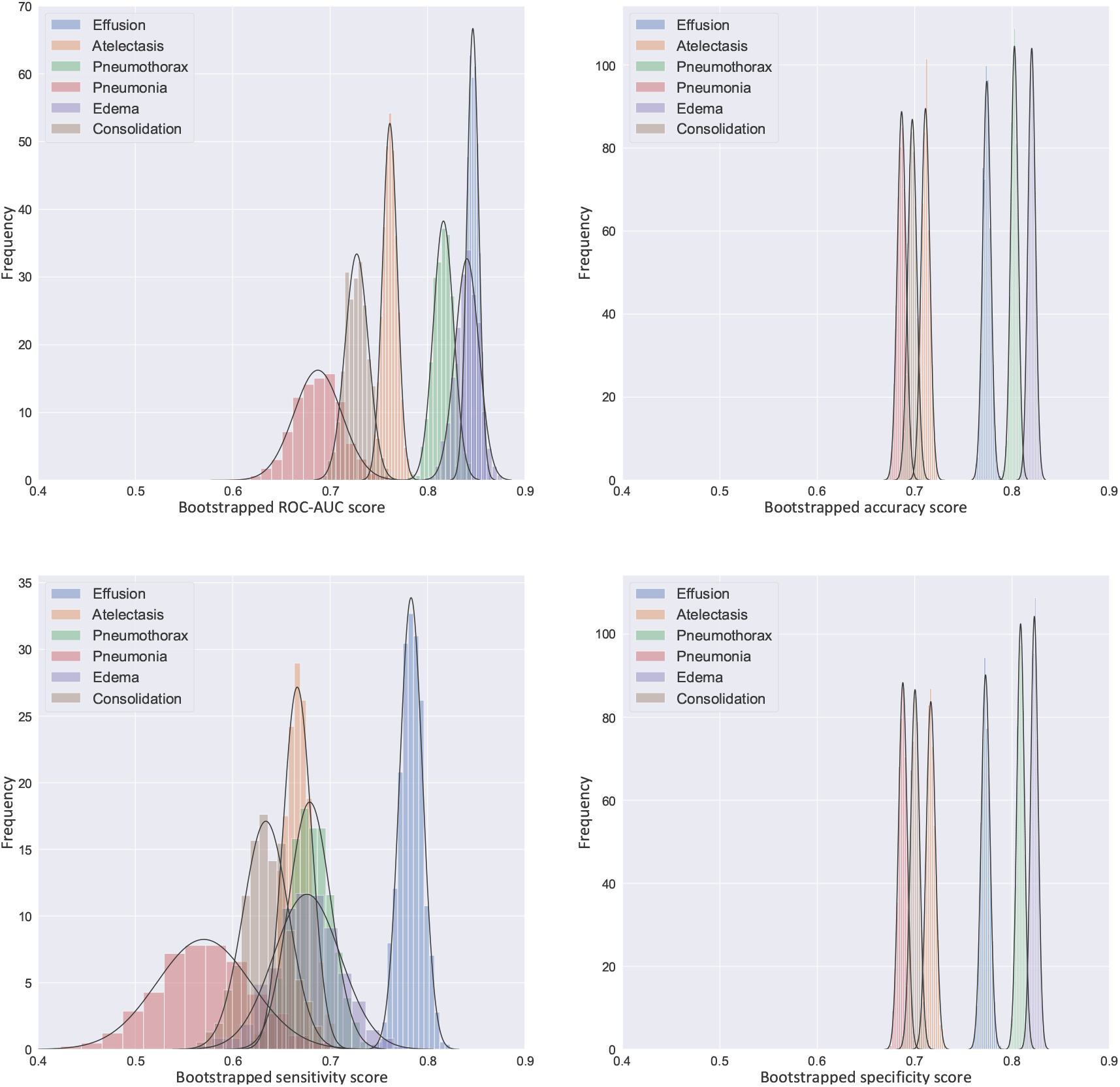
Distribution of performance measures when analysed via bootstrapping. ROC-AUC, accuracy, sensitivity, and specificity were selected as performance measures in the data merging (NIH + Stanford GAN) experiment. In boostrapping experiments, we randomly resampled 10,000 bootstrap replications from the test set (22,424 radiographs) to obtain the statistical distribution of the found values. The means and the standard deviations of the selected performance measures were estimated by Gaussian fitting (black curves). Abbreviations: ROC-AUC, area under receiver operating characteristic curve.

